# The need for unrealistic experiments in global change biology

**DOI:** 10.1101/2021.12.07.471575

**Authors:** Sinéad Collins, Mridul K. Thomas

## Abstract

Climate change is an existential threat, and our ability to conduct experiments on how organisms will respond to it is limited by logistics and resources, making it vital that experiments be maximally useful. The majority of experiments on phytoplankton responses to warming and CO_2_ use only two levels of each driver. However, to project the characters of future populations, we need a mechanistic and generalizable explanation for how phytoplankton respond to concurrent changes in temperature and CO_2_. This requires experiments with more driver levels, to produce response surfaces that can aid in the development of predictive models. We recommend prioritizing experiments or programmes that produce such response surfaces on multiple scales for phytoplankton.

---

Biological research today is dominated by anthropogenic climate change and the existential threats it poses. Biologists are therefore focused on understanding how rapid environmental change affects organisms, populations, communities and ecosystems, on spatial scales ranging from local to global, and temporal scales ranging from seconds to decades. As in other areas of science, this research is divided into fundamental and applied. Applied research has straightforward practical justifications: we need to improve food production, conservation, predict and mitigate the spread of infections, effectively protect ecosystems to preserve key services, or project how continued human activities will impact other species. However, the majority of research on biological responses to climate change falls under the banner of fundamental science, which aims to provide insights, examples, and theoretical frameworks that help move knowledge (including applied science) forward.

Here we consider the fit-to-purpose of experiments designed to investigate organismal responses to environmental change. Our aim is to evaluate fundamental experiments that use two or more simultaneous environmental changes, which are often called ‘multiple stressor’, ‘multi-stressor’ or ‘multiple driver’ experiments in the aquatic literature [1]. We focus on experiments investigating the joint effects of changing CO_2_ and temperature levels as a case study. We look at how their design helps us make progress towards projecting how they jointly affect phytoplankton traits underlying ecological or biogeochemical function in aquatic systems. We highlight how fundamental research can improve our understanding of biotic responses to complex environmental change by expanding experimental designs to include unrealistic environmental conditions that no foreseeable future holds.

## BOX 1

**Trait:** A measurable quantity describing a particular organismal or population function/quantity, usually related to organismal fitness, ecological or biogeochemical function (e.g. population growth rate, cell size, nutrient affinities). For a more detailed discussion of phytoplankton trait types see [2]

**Response curve/reaction norm:** The *shape* of the biological response to a change in some continuous variable (e.g. temperature response curve). See figures 2-4.

**Response/interaction surface:** The shape of the surface that describes the biological response to joint changes in two or more continuous variables (e.g. growth responses to concurrent changes in CO_2_ and temperature). See figures 2-4.

**Experiment:** We focus here on manipulative studies carried out in controlled environments, usually in laboratories, where traits are measured and trait changes can be unambiguously linked to specific environmental changes.

END BOX 1

## What ocean global change questions do we want to answer with laboratory experiments?

The goals of ocean global change biology are clear. Among them are predictions of how ecosystems and biogeochemical cycles are likely to change globally that are good enough to take effective action or at least anticipate large-scale changes [3]. Fundamental research is aimed at understanding how the environment affects (and is affected by) biota, and at projecting future dynamics and distributions of species, communities, ecosystems, ecosystem services, and biogeochemical cycles. The questions typically asked in global change biology can be summarised as: *what does the future bring at X organisational scale and Y geographic scale*? To project change in specific ecosystems, for specific timescales and projections of environmental change, these questions are made concrete. For primary producers in aquatic systems, such questions include: How much and where will primary production change as a result of expected changes in CO_2_, temperature, and nutrient levels? How will warming change the biological carbon pump in the open ocean? How will nitrogen input into the oceans change as warming increases the range of N-fixers?

Connecting these large-scale questions to the ecophysiological processes that underpin them partly boils down to understanding the environmental dependence of key organismal traits and processes: population growth rates, cell compositions, resource uptake rates, edibility, cell sizes and more. Understanding how the environmental dependence of these traits will shape future ecosystem processes requires integrating and modelling ecophysiology, community ecology, and evolution. The traits that need to be understood in phytoplankton are largely known and measurable in the laboratory [4], and there is sufficient understanding of the physiology underlying these traits to conduct hypothesis-driven experiments on mechanisms underlying known or putative tradeoffs between traits in many cases [5]. At this point, the major empirical knowledge gap is robustly connecting variation in trait values to variation in the environment. This is a tractable problem that can be addressed with experiments.

Dynamic models can simulate the growth of species (or higher taxonomic units) alone or in mixed communities, with and without predators, at temperatures or CO_2_ levels that were not measured (extrapolating requires caution, however), in constant or variable environments, and many can account for eco-evolutionary processes [6-9]. These involve important assumptions, but represent vital steps towards the larger goal of making accurate predictions of future populations and ecosystem change. To model how populations and communities will change when both temperature and CO_2_ change, these dynamic models need parameterised equations that capture the temperature-CO_2_ *response surface*. At present, we do not have the data to tell us the shape of this surface, so we lack an equation to describe it (see Figs. 2-4 for one possibility, based on [10]. However, this can be solved with experiments that we have the methods to do, but rarely carry out. To produce response surfaces, it can be unhelpful to replicate only realistic environmental conditions. Equations for temperature-CO_2_ response surfaces would be constrained better if experiments included high temperature-low CO_2_ treatments, and vice versa.

## How are we studying driver interactions at present?

We compiled data on temperature x CO_2_ interaction experiments involving two key phytoplankton taxa (diatoms and coccolithophores). We included data from studies that manipulated both CO_2_ and temperature, with at minimum a 2 × 2 design, and which measured population growth rate. We found 16 studies that met these criteria (see [11,12] for a representative example). Nearly all the data from these studies have been gathered at just 2 levels of CO_2_ (∼400ppm and ∼1000ppm), at or near 20°C (Fig. 1). Despite omitting simple cases where drivers were not independently manipulated, the median experiment is still a 2 × 2 study. Overall, the published data do a poor job of sampling the interaction surface for temperature and CO_2_. This is concerning because while we can (and do) produce explanations and models based on basic physiology and conserved tradeoff phenomena that are consistent with the variation in outcomes in the published data [13-15], our current understanding of physiology cannot build interaction surfaces from first principles.

**Fig. 1.**
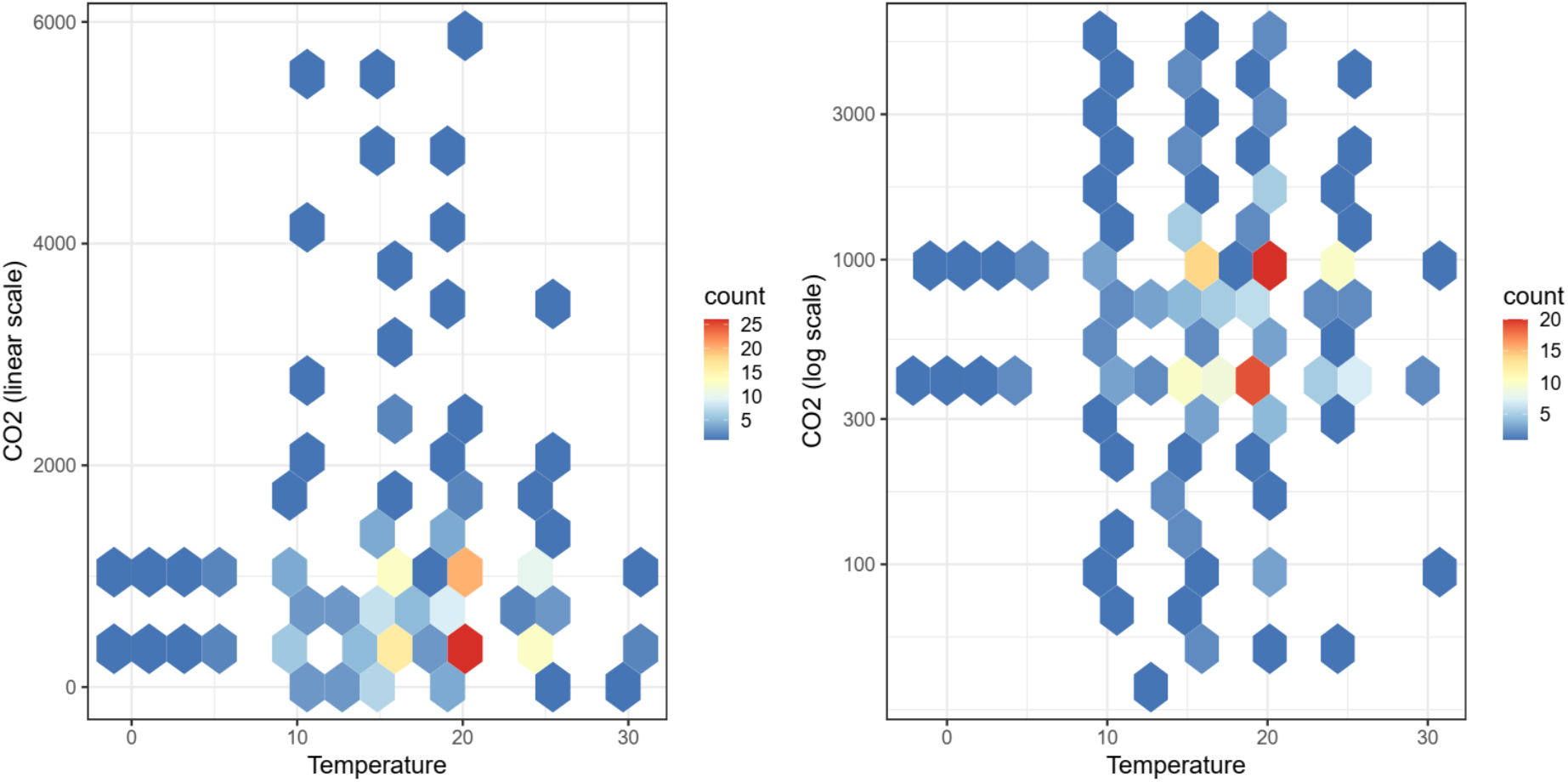
**What temperature and CO_2_ values are we measuring? The colour of each hexagon indicates the number of measurements in our compiled dataset at (narrowly binned) combinations of temperature and CO_2_ values. Note that different studies have different numbers of measurements. Even allowing for the fact that very high concentrations are of less interest for forecasting future responses, there are large gaps in the data we collect. Tropical and polar temperatures are underrepresented, as well as CO_2_ levels between 400 and 1000 ppm. Data are from single-genotype laboratory experiments with diatoms and coccolithophores where at least 2 levels of CO_2_ and 2 levels of temperature were used, and growth rates were reported [11,16-31]. See Supplementary information for details of search and data.**

**Fig. 2.**
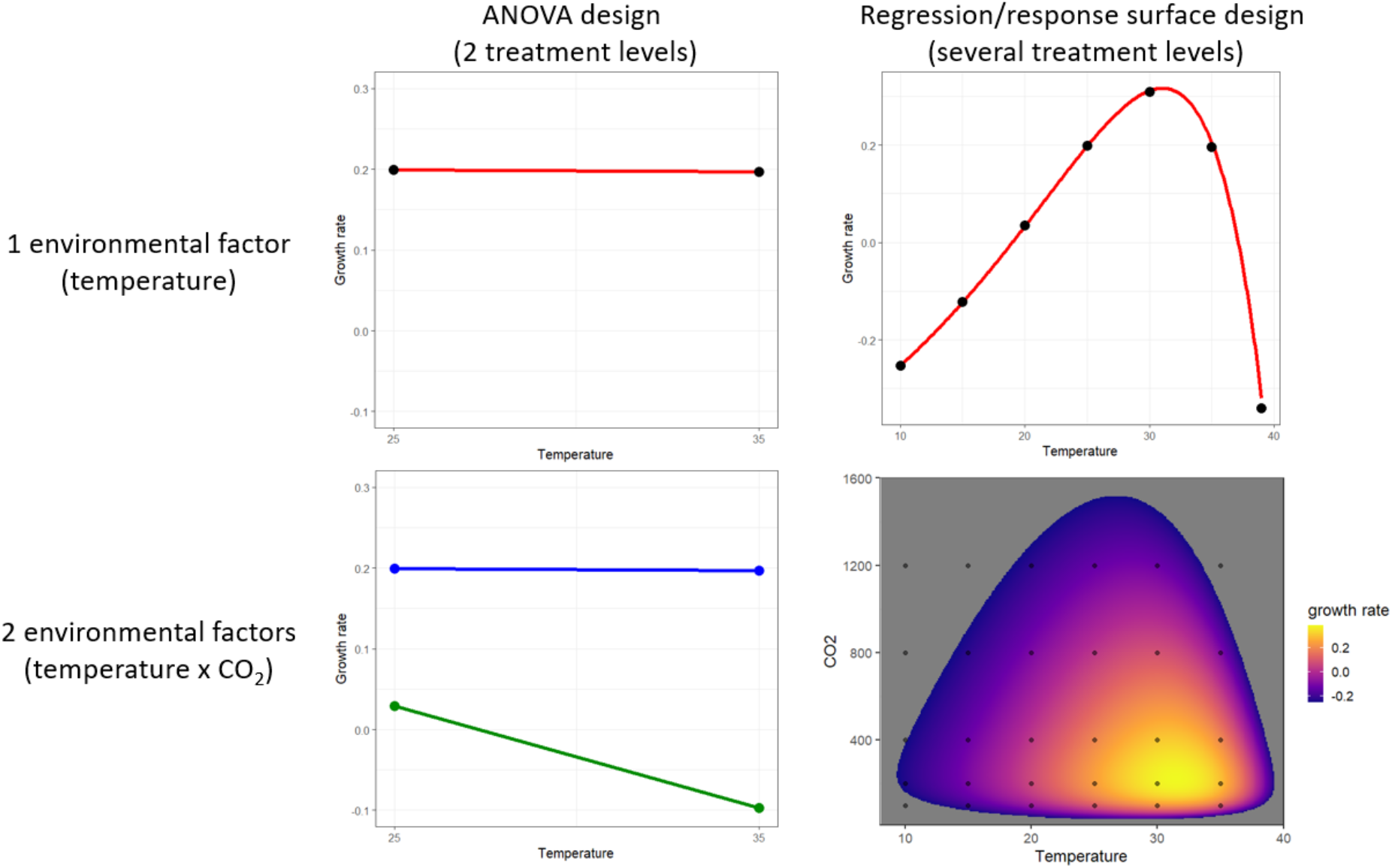
**Experimental design for continuous variables like temperature and CO_2_. Present experiments treat both temperature and CO_2_ as discrete and measure them at 2 levels (left panels). This limits our understanding, inferences and ability to build on experiments with modelling efforts because in reality, responses to both are continuous and nonlinear. The response to temperature is left-skewed and unimodal (top right). The response to CO_2_is right-skewed and unimodal (not shown). Jointly, they form a complex response surface that we do not understand well at present for lack of data. Bottom right shows a hypothetical surface based on a temperature × CO_2_model described in the supplementary information. Growth rates below -0.25 are suppressed (dark grey) to highlight variation in positive growth rates.**

## Key limitations of the current approach to driver interaction experiments

Temperature response curves exist for >100 species representing all major phytoplankton functional groups [32]. CO_2_ response curves at different temperatures are scarce, so expectations of temperature-CO_2_ response surfaces cannot be generated. Our estimates of organismal responses to the joint effects of rising CO_2_ and warming are therefore sensitive to small errors in both predictions of future environments (Figure 3) and assumptions about the shape of the response surface (Fig. 4). Despite the knowledge that warming and changes to CO_2_ are key drivers in marine systems, and decades of experiments with temperature and CO_2_, we have yet to build robust hypotheses of how even model phytoplankton should respond to the joint action of these drivers.

**Fig. 3.**
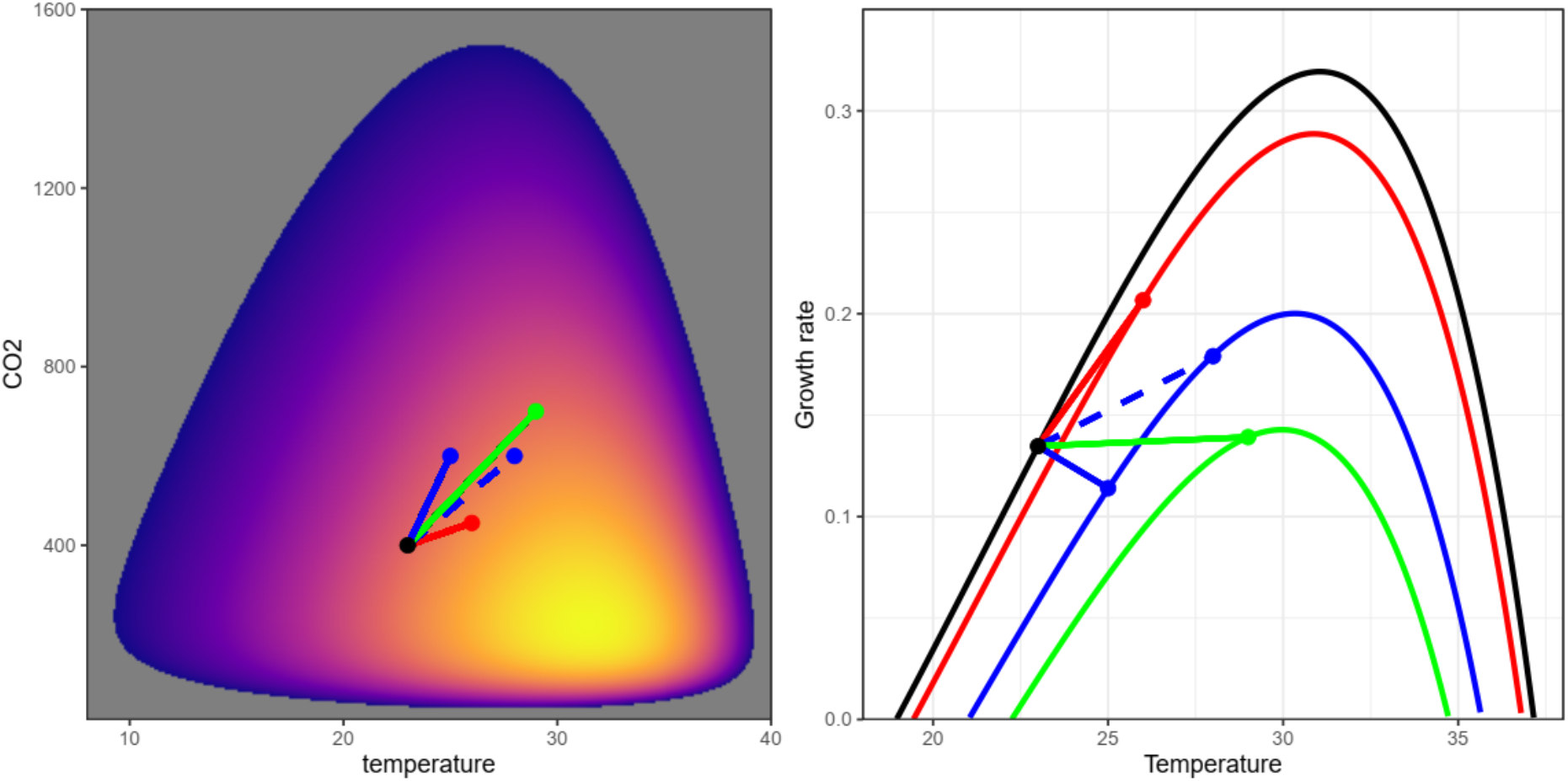
**Uncertainty in both temperature and future CO_2_ is hard to account for in ANOVA designs. Growth rates in present day conditions (represented by black point) may increase, decrease or stay substantially the same based on small differences in future conditions (represented by coloured points). Coloured curves in the right panel represent temperature curves at different CO_2_ concentrations i.e. horizontal slices through the response surface in the left panel. Conclusions drawn from existing 2 × 2 experiments are therefore heavily dependent on the reliability of CO_2_ projections - which themselves depend on the phytoplankton response - and climate sensitivity (compare blue points, which represent different amounts of warming based on the same amount of CO_2_). We can escape this problem of circularity only by understanding the response surface better.**

**Fig. 4.**
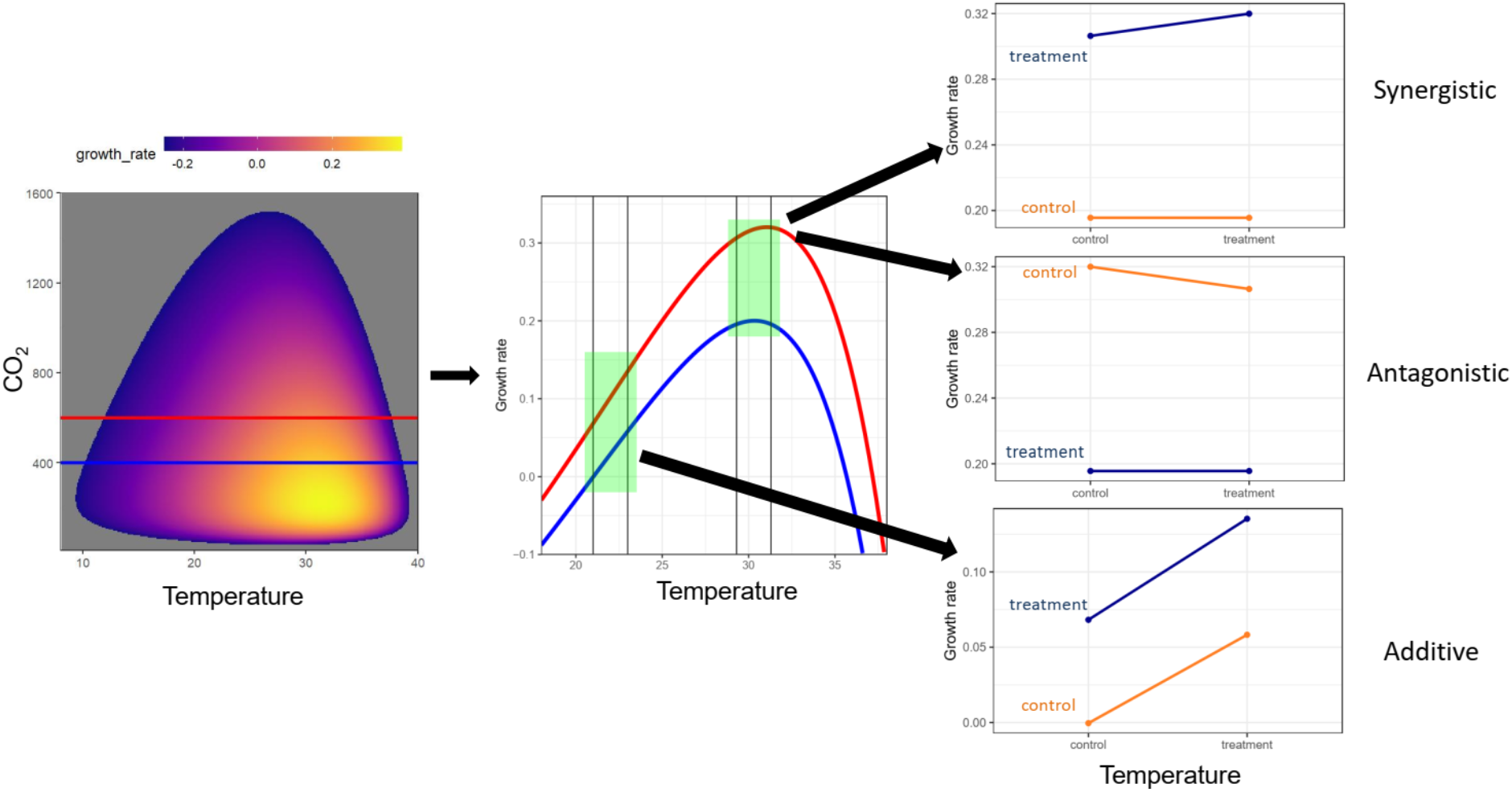
**Response surface experiments can lead to very different inferences than ANOVA experiments. In this example, the same response surface (left panel) is reduced first to 2 response curves (middle panel) and finally to 2 × 2 ANOVA experiments (right panels). The specific ANOVA experimental levels chosen (vertical black lines in middle panel) and the choice of reference level alters the nature of the inferred interaction. Note that the synergistic and antagonistic interactions shown are from the same set of 4 growth rates. In the synergistic case, the low CO_2_, low temperature point is chosen as control, while in the antagonistic case, the high CO_2_, high temperature point is chosen as control. Additive/synergistic/antagonistic interactions are context-dependent, whereas capturing the response surface largely eliminates the need for this misleading taxonomy.**

Our compilation shows that the literature is dominated by ANOVA experimental designs, which are appropriate for discrete variables (e.g. species A vs species B). These designs treat continuous variables as discrete (e.g. temperature treated as ‘present’ vs ‘future’). In contrast, regression designs appropriate for continuous variables involve using many experimental levels of the variable and modelling a continuous response. Response surface designs are multi-dimensional versions of regression designs (Fig. 2).

When would 2 × 2 ANOVA experiments achieve the goal of understanding and predicting how projected environmental change affects organisms? If we had precise and accurate forecasts of temperature and CO_2_ levels, it could be argued that predicting changes in a species’ performance only requires comparing performance under present and perfectly-predicted future conditions. However, we do not know precisely what CO_2_ levels or even average global temperatures will be decades from now, with uncertainties running to several degrees [33]. Simulations from a realistic response surface (Fig. 3) show that modest uncertainty in either dimension can lead to substantially different outcomes. This uncertainty is amplified by variation through time and across space, and in other environmental dimensions. Organismal responses also vary based on evolutionary differences and ecological interactions. This variation in outcomes does not mean experiments measuring biological responses to the environment are fruitless. Uncertainty in biological responses can be accounted for if we measure and model responses appropriately. An ANOVA-based approach cannot account for most of this uncertainty. Indeed, it is difficult from the current literature to partition how much of the observed variation in responses to combinations of ocean acidification and warming in phytoplankton [13,34] is attributable to biological variation in responses, and how much is due to uncertainty introduced by small differences in the way experiments sample interaction surfaces.

The ANOVA approach is not rescued by binning responses into qualitative levels. The common additive/synergistic/antagonistic interaction framework and its extensions or variations [35,36] is subject to the same weaknesses described above, and additional ones too. Differences in the precise driver levels chosen, and the choice of the control/reference level can lead to any of the three possible outcomes from the same response surface (Fig. 4). The additive/ synergistic/ antagonistic framework is often applied to higher levels of organisation such as ecosystem responses and it is possible that some of our objections do not apply there; this merits more investigation than it has received.

ANOVA-based approaches also make it challenging to draw conclusions about responses at higher taxonomic levels without additional experiments targeting those levels. In contrast, response surfaces allow scaling across levels more easily. The ‘Eppley Curve’ is a classic oceanography community pattern that arises from species temperature response curves [37,38]. It defines an upper boundary on maximum growth rate that increases exponentially with temperature and can be constructed as the ‘envelope’ of a series of temperature response curves. This connection between an important community-level pattern (the Eppley curve) and population-level response curves allows for easy switching between scales of interest - a clear advantage for modelling and prediction. A multi-dimensional version of the Eppley curve that varies with CO_2_ concentration (an Eppley surface) should be possible with sufficient data on variation in temperature-CO_2_ response surfaces.

## Conclusions

To have a robust understanding of how phytoplankton (and other organisms) respond to multiple simultaneous drivers, we need to move beyond ANOVA-based experiments that categorise interactions as additive, synergistic, antagonistic (or variations thereof), and towards experiments that produce interaction/response surfaces. Regression designs are logistically more challenging and often require better experimental facilities than ANOVA-based ones. The nonlinearity of the temperature response curve is not captured well with fewer than 5 temperature levels; more is better. Based on data thus far, the same appears to be true for CO_2_ response curves [10]. ‘Collapsed designs’ can be useful in reducing the size of experiments relative to fully-factorial designs (see [39]) and the more information we have about the shape of the surface, the better equipped we will be to intelligently sample it and parameterise it for a species. However, collapsed designs do not replace fully factorial experiments, which are often doable for microbes, given adequate resourcing and prioritization. For example, a 6 × 6 experiment can involve just 36 experimental units (replication is not necessary for regression designs though it can be helpful if specific levels are particularly important for parameter estimation). Depending on the traits under investigation, this is a challenging but doable experiment. Indeed, both Sett (2014) [19] and Zhang (2020) [25] conducted experiments with enough total experimental units to have calculated rough response surfaces had more levels of temperature been used; both used more CO_2_ levels than necessary.

There is substantial variation in responses to changes in temperature and CO_2_ both within and between functional groups (and within and between species) [10,32,34,40,41]. While studying response surfaces for model organisms from each functional group will produce general insights, estimates of how much taxonomic variation exists for response surfaces are also needed. Replicate experiments are necessary in order to estimate the extent of biological variation in the biological responses that form interaction surfaces; repeating experiments, either within single studies that encompass multiple genotypes or species (e.g. [40]), or as coordinated networks of experiments (e.g.[42]) is both useful and necessary; funding and publication success must begin to reflect this. Finally, phytoplankton have the capacity to evolve between now and nearly any future date we make projections for, and evolutionary responses can differ from and even reverse plastic ones [43-45]. There is also good evidence that adaptation to warming is affected by other drivers [46]. We need generalizable insights into how response surfaces evolve under different rates and patterns of multidriver environmental change.

We suggest prioritizing the measurement of response surfaces at multiple timescales to capture plastic/acclimatory as well as eco-evolutionary responses for phytoplankton that can be grown in the laboratory. This calls for allocating resources to larger or coordinated experiments. We recognize this is challenging given funding cycles and the need to publish frequently, both of which favour smaller, shorter, stand-alone experiments. However, producing data that reveals response surfaces is what basic science is meant to do: lead to usable insights about real-world phenomena and inspire theoretical frameworks that help us understand and predict the world better. We think that this is a better investment of limited resources than the continued production of smaller or isolated experiments focused almost entirely on realistic environments that, from our current position, have made limited headway towards a usable understanding of phytoplankton responses to multidriver environmental change.

## Supporting information

Supplemental materia

Data for Figure 1

## Acknowledgments

We thank Allanah Paul and Lennart Bach for discussions of their model. The work presented in this article results, in part, from funding provided by national committees of the Scientific Committee on Oceanic Research (SCOR) and from a grant to SCOR from the U.S. National Science Foundation (OCE-1840868) to the Changing Oceans Biological Systems project, and in part from funding to SC from the Gordon and Betty Moore Foundation (MMI 7397).

