## Supplemental materia for "The need for unrealistic experiments in global change biology"

### Supplementary Information

### I. Model description

To create a new model describing how temperature and CO2 interact to affect phytoplankton growth, we adapted two existing models, one of growth depending on both temperature and nutrients (Thomas et al. 2017) and the other of growth depending on $CO_2$ (or more precisely, $[H^+]$, Paul & Bach 2020). This new temperature x CO2 model is intended to be illustrative, not a solution to the problems we describe in the main text. We still need to develop an accurate, predictive model of temperature-$CO_2$ interactions, and this will require more and better experimental data to achieve.

*1. Modelling how growth depends on temperature*

From equation 1 in Thomas et al. (2017), we extracted the temperature-dependence of growth, shown below:

$$\mu(T) = b_1 \cdot \exp(b_2 \cdot T) - (d_0 + d_1 \cdot \exp(d_2 \cdot T)) \tag{1}$$

Where specific growth rate $\mu$ depends on temperature $T$. $b_1$ is the birth rate at a temperature of 0°C, $b_2$ is the exponential change in birth rate with increasing temperature, $d_0$ is a temperature-independent mortality term, and $d_1$ and $d_2$ jointly describe the exponential increase in mortality rate with temperature (see Thomas et al. 2017 for further details).

*2. Modelling how growth depends on $CO_2$*

Paul & Bach (2020) proposed that a single master equation governs the growth of all phytoplankton as a function of CO2, with concentrations below a threshold being limiting and above it being inhibiting. This equation (Equation 1 in Paul & Bach 2020) describes how growth depends on $[H^+]$ and not $CO_2$ directly:

$$\mu([H^+]) = \mu_{max} \cdot \left(1 - \frac{[H^+]}{[H^+]^*}\right)^n \cdot \left(\frac{[H^+]}{[H^+] + C_m\left(1 - \frac{[H^+]}{[H^+]^*}\right)^m}\right) \tag{2}$$

where $\mu_{max}$ is the maximum growth rate, $[H^+]$ is the proton concentration, $[H^+]^*$ is the critical inhibitor concentration above which no growth occurs, $C_m$ is the Monod constant, and $n$ & $m$ are constants (see Paul & Bach 2020 for further details). We omit one scaling term from Paul & Bach 2020, *[cell]*, in order to model the per capita population growth rate.

Note that the equation is parameterised in terms of $[H^+]$ and not $CO_2$. Converting between $[H^+]$ and $CO_2$ is not straightforward because of the complexities of carbonate chemistry and the authors made this conversion using the R package *seacarb* (Gattuso et al. 2021). This involves making some simplifying assumptions (e.g. constant total alkalinity) but appears successful in describing empirical variation in $CO_2$-dependent growth.

*3. Modelling how growth depends on the joint effects of temperature and $CO_2$*

We unite these two models below by proposing that the $CO_2$-dependent growth in Paul & Bach (2020) may be multiplied by the increasing ('birth') part of the temperature-dependent growth equation. Temperature-dependent death rates are then subtracted from the birth part of the equation.

$$\mu(T,[H^+]) = \left[b_1 \cdot \exp(b_2 \cdot T) \cdot \mu_{max} \cdot \left(1 - \frac{[H^+]}{[H^+]^*}\right)^n \cdot \left(\frac{[H^+]}{[H^+] + C_m\left(1 - \frac{[H^+]}{[H^+]^*}\right)^m}\right)\right] - (d_0 + d_1 \cdot \exp(d_2 \cdot T)) \quad (3)$$

We follow Paul & Bach's step of converting between $[H^+]$ and $CO_2$ using the R package *seacarb* (Gattuso et al. 2021) and assuming constant total alkalinity in the conversion, thereby allowing us to model not just $\mu(T,[H^+])$ but $\mu(T,CO_2)$. We do account for the effects of temperature while making the conversion using *seacarb*.

Equation (3) can be simplified further, for example by considering the $\mu_{max}$ term. But the equation itself is simplistic and we do not expect it to accurately capture all important aspects of temperature-$CO_2$ interactions. It serves here to illustrate the value of being able to model a temperature x $CO_2$ response surface. It also highlights the fact that we are desperately lacking in data with which to make such models; we presently do not have the data we would need to identify the model's flaws and refine (or reject) it.

### II. Model parameterisation

The parameters we used for the response surface plots in the main text are shown below:

| Parameter name | Parameter value |
|---|---|
| b1 | 0.300198208 |
| b2 | 0.057696903 |
| d0 | 0.679246924 |
| d1 | 0.001869989 |
| d2 | 0.178471387 |
| $\mu_{max}$ | 1 |
| $[H^+]^*$ | $6.50 \times 10^{-8}$ |
| $C_m$ | $4.31 \times 10^{-9}$ |
| n | 1.31 |
| m | 33.1 |

The $CO_2$-dependent growth parameters (except for $\mu_{max}$) were taken from the 'heavily calcified coccolithophores' parameter values in Paul & Bach (2020). $\mu_{max}$ was set at 1 for simplicity, and the temperature-dependent parameters were chosen for convenience and do not reflect any specific species.

### III. Data for Figure 1

Google Scholar was used to search for articles in January 2021. Using the search term 'temp* OR warm*' AND "acidification" OR "pH" OR "CO2" OR "carbon dioxide" AND 'diatom* OR coccolithophore* OR "phytoplankton" AND "driver" OR "interact"' AND '-review'. This returned 186 papers after removing duplicates. Studies were excluded based on the following criteria: No primary data, no growth rates reported, study organism not a coccolithophore or diatom, warming and $CO_2$ not manipulated independently, warming and acidification not combined, author retraction, data not available with no author response when asked to provide data, confounding variables not controlled, no 'control' or reference conditions, experiment has no replication, no sample size reported, no standard deviation reported. This resulted in a final count of 16 publications. For publications containing data on more than one genotype/strain, each genotype/strain is counted as a single "study" in Figure 1. See Supplementary Data excel sheet for details of each study.
